# Response to early drought stress and identification of QTLs controlling biomass production under drought in pearl millet

**DOI:** 10.1101/373233

**Authors:** M Debieu, B Sine, S Passot, A Grondin, AE Akata, P Gangashetty, V Vadez, P Gantet, D Foncéka, L Cournac, CT Hash, NA Kane, Y Vigouroux, L Laplaze

## Abstract

Pearl millet plays a major role in food security in arid and semi-arid areas of Africa and India. However, it lags behind the other cereal crops in terms of genetic improvement. The recent sequencing of its genome opens the way to the use of modern genomic tools for breeding. Our study aimed at identifying genetic components involved in early drought stress tolerance as a first step toward the development of improved pearl millet varieties or hybrids. A panel of 188 inbred lines from West Africa was phenotyped under early drought stress and well-irrigated conditions. We found a strong impact of drought stress on yield components. This impact was variable between inbred lines. We then performed an association analysis with a total of 392,493 SNPs identified using Genotyping-by-Sequencing (GBS). Correcting for genetic relatedness, genome wide association study identified QTLs for biomass production in early drought stress conditions and for stay-green trait. In particular, genes involved in the sirohaem and wax biosynthesis pathways were found to co-locate with association loci. Our results open the way to use genomic selection to breed pearl millet lines with improved yield under drought stress.

## Introduction

Pearl millet (*Pennisetum glaucum* (L.) R. Br., syn. *Cenchrus americanus* (L.) Morrone) was domesticated in the central Sahel region of Africa [1]. It is adapted to dry and hot climates and it plays a major role in food security in arid and semi-arid areas of Africa and India. However, there is still a wide scope to increase its adaptation to environmental constraints limiting its production. Pearl millet is the staple crop of more than 90 million farmers. Grains are gluten-free and rich in proteins and essential micronutrients such as iron or zinc. It is consumed directly as couscous or porridge and also used to make flour to produce flat bread or to be introduced in bread production to reduce the imports of wheat in some African countries. Moreover, the aerial biomass is an important source of fodder for animals. However, pearl millet lags behind the other cereal crops in terms of genetic improvement and its average yields are still low. The recent sequencing of its genome [2] opens the way to tap on its large genetic diversity in pearl millet to breed varieties and hybrids adapted to current and predicted future climatic constraints [3].

Drought is one of the major factors limiting pearl millet production. In sub-Saharan Africa, pearl millet is mostly grown in areas characterized by low rainfall and sandy soils with very little organic matter that have therefore low water retention ability. The climate is characterized by a long dry season and a short rainy season where most of the rain-fed agriculture is concentrated [4]. Pearl millet is usually sown after or just before the first rain of the rainy season. Because of the rain pattern, pearl millet can face drought stress at early stages if the first rains of the season are distant from each other. Another important drought stress period is the grain filling stage. Current climate models predict that the rain pattern in sub-Saharan Africa will be more variable from one year to the other and will involve more extreme events [4]. Episodes of drought are therefore expected to be more frequent during the rainy season. Altogether, climate change is expected to lead to a reduction in pearl millet yield in the area [4]. While terminal drought stress was the focus of several studies in pearl millet [5], [6], [7], [8] little is known about the impact of drought episodes during the vegetative phase.

The aim of this study was to characterize the impact of early drought episodes on pearl millet in field conditions and to identify genetic components contributing to tolerance to this stress. A panel of 188 inbred lines was grown during two successive seasons under well watered and early drought stress conditions in Bambey (Senegal) and was phenotyped for several agromorphological traits. We found a strong impact of early drought stress on yield. Genotyping by sequencing (GBS) data were generated and allowed the identification of 392493 SNPs. Genome wide association study (GWAS) using a mixed model and correcting for genetic relatedness identified potential quantitative trait loci (QTLs) for biomass under early drought stress and stay-green character.

## Material and Methods

### Plant material and trait measurements

A panel of 188 Pearl millet (*Pennisetum glaucum (L.) R. Br.*) inbred lines developed at ICRISAT (Niger) from landrace and improved open-pollinated cultivars of West African origin was used in this study.

Field trials were performed at the CNRA (Centre National de Recherche Agronomique) in Bambey (14.42°N, 16.28°W) in Senegal, in February 2015 and 2016 during the dry season on two adjacent fields. Soils are deep sandy soil with low levels of clay and silt (12%) and organic matter (0.4%). Clay and silt content increase with soil depth from 10.2% in the 0 to 0.2 m layer to 13.3% in the 0.8 to 1.2 m layer. Trials were set up using an incomplete randomized blocks design (S1 & S2 Figs). A total of 20 plants were grown per variety in 2 rows of 3 m long with 0.30 m between plants and 0.8 m between rows. Fifteen days after sowing, a single plant per planting hole was conserved, so ten plants per variety for a given row. Fertilization (NPK) followed standard recommendation i.e. 150 kg ha^−1^ of NPK (15-15-15) after sowing. Urea was applied at 100 kg ha^−1^ at two stages, 50 kg ha^−1^ after thinning and 50 kg ha^−1^ 30 days after sowing.

The flowering date corresponding to 50% of plants flowering (in days after sowing) was recorded. Height of the plant (in cm), panicle and stalk length (in cm), total number of leaves, aerial biomass (Kg.plant^−1^), as well as stay-green trait (number of green leaves/total number of leaves) were measured on 6 plants per plot at maturity. Shoots and panicles were harvested and used to quantify total grain weight (g/plant), number of grains per plant, weight of 1000 grains (g) and aerial biomass (g dry mass/plant). These data were used to compute a harvest index (grain weight/biomass weight).

Plants were grown during the dry season (February to June) to fully control the amount of water provided (no rain during the experiments). Irrigation was performed twice per week with 30 mm water per irrigation. Early drought stress was applied by stopping irrigation 3 weeks after germination for 4 weeks. Irrigation was then resumed until the end of the growth cycle. Field dry-down was followed by measuring volumetric soil moisture to evaluate the fraction of transpirable soil water (FTSW) using Diviner probes (Sentek Pty Ltd). Canopy temperature was measured on 2 plants per plot 30 days after the start of the dry down period in both well-irrigated and drought-stress treatments using an Infrared thermometer (Quicktemp).

The spatial trend in the incomplete block design was evaluated using control plants (inbred line ICMB88004) and the phenotype of each line was corrected using the SpATS package in R (S3 Fig; [9]). Corrected data are provided as S1 Data.

### DNA isolation and genotyping by sequencing

Genomic DNA was isolated from three-week old leaves as previously described [10]. A single plant was used per accession and 68 were duplicated as controls. Whole-genome genotyping was carried out using Genotyping-By-Sequencing (GBS) at the Genomic Diversity Foundation at Cornell University (Ithaca, USA). GBS libraries were prepared using restriction enzyme *Ape*KI [11]. Digested DNAs were ligated to barcoded adapter pairs. 96- plex libraries were sequenced using a HiSeq2500 and a NextSeq500 sequencing system (Illumina). Adapter sequences were removed using the cutadapt software and low quality reads were filtered out with the pearl script Filter_Fastq_On_Mean_Quality.pl from SouthGreenPlatform (Minimum mean quality allowed for a read=30, Minimum length allowed for a read=35). The reads were mapped using BWA [12], [13]. Picard-tools-1.119 and Genome Analysis ToolKit (GATK-3.6 algorithm UnifiedGenotyper) softwares were used to create the vcf file. SNPs were filtered out based on missingness percentage (<50% missingness), homozygosity (inbreeding F>0.5) and minor allele frequency (MAF ≥ 5%).

### Population structure analysis

A neighbor joining (NJ) phylogenetic tree was generated with TASSEL v5.2.39. Population structure was assessed with a random subset, with one SNP/100kb using the program sNMF. Five runs were performed for a given number of group (K) and the values with the smallest Cross-Entropy for each K were selected to generate the structure graph.

### Genome wide association study (GWAS)

Marker-trait associations were established using pearl millet inbred lines phenotyped for 11 agromorphological traits under two conditions (early drought stress and well-watered) and in 2 years (2015 and 2016). For each year, we performed GWAS with a mixed model correcting for genetic relatedness (kinship matrix) using the R package GAPIT (Genome Association and Prediction Integrated Tool; [14], [15]). The *p*-values obtained for both years were combined using a Fisher combining probability method [16]. We considered a *False Discovery Rate* (*FDR*, threshold = 0.1) in determining the significant SNPs.

## Results

### Phenotypic variation under normal and early drought stress conditions

In order to evaluate pearl millet response to early drought stress, a panel of 188 inbred lines was characterized on well watered (WW) and early drought stress (DS) conditions on two successive years in Bambey (Senegal). In the well watered treatment, the fraction of transpirable soil water (FTSW) was generally above 40% along the soil profile (0-120 cm) across the experiment in both years, indicating that water was not limiting for plant growth (S4 Fig; [17]). In drought stress treatments, FTSW was below 40% in 0-60 cm soil profiles at 40 days after sowing (DAS) in 2015 and at 49 days in 2016. This measure indicated efficient field dry-down and imposition of water limiting conditions at vegetative stage (S4 Fig). In 2015, water remained limiting for plant growth (FTSW below 40%) along the soil profile (0-120 cm) until around 110 DAS, extending the water-limiting period to the reproductive stage. In 2016, irrigation of the field at 49 DAS allowed increase in FTSW to around 40% at 55 DAS between 60-120 cm, although short periods of water limiting conditions appeared at reproductive stage (around 75 and 85 DAS). In 2015 and 2016, canopy temperature measured 30 days after the start of the dry-down was significantly increased in the drought-stress treatment as compared to the well-watered treatment (ANOVA, p < 0.001), indicating that plants were indeed subjected to drought stress (S5 Fig).

In order to evaluate the impact of early drought stress, a total of 11 agromorphological traits were measured at the seed maturation stage (Table 1). For most of the traits, a large range of variation was detected, with the coefficients of variation (CV) varying from 0.082 for stay-green under well watered conditions to 0.986 for total grain weight under drought stress conditions. Most of the traits showed a normal distribution (S6 Fig) apart from flowering date that was flatter than a normal distribution (negative kurtosis). For each trait, each year and all conditions, variance analyses showed a strong significant genetic effect on phenotypic variation (Table 2). A high correlation was found between results obtained in the two successive years both for well watered (WW) and drought stress (DS) conditions indicating that the response was comparable in both years (Fig 1). Analyses of relation between traits showed that some traits were highly correlated (Fig 2; S1 Table) such as total grain number and total grain weight. Correlated traits under drought stress conditions were generally also correlated under well watered conditions (correlation of correlation coefficients, *r* > 0.8, *p*<10^−12^, Figure 2 C D; S1 Table).

**Table 1.**
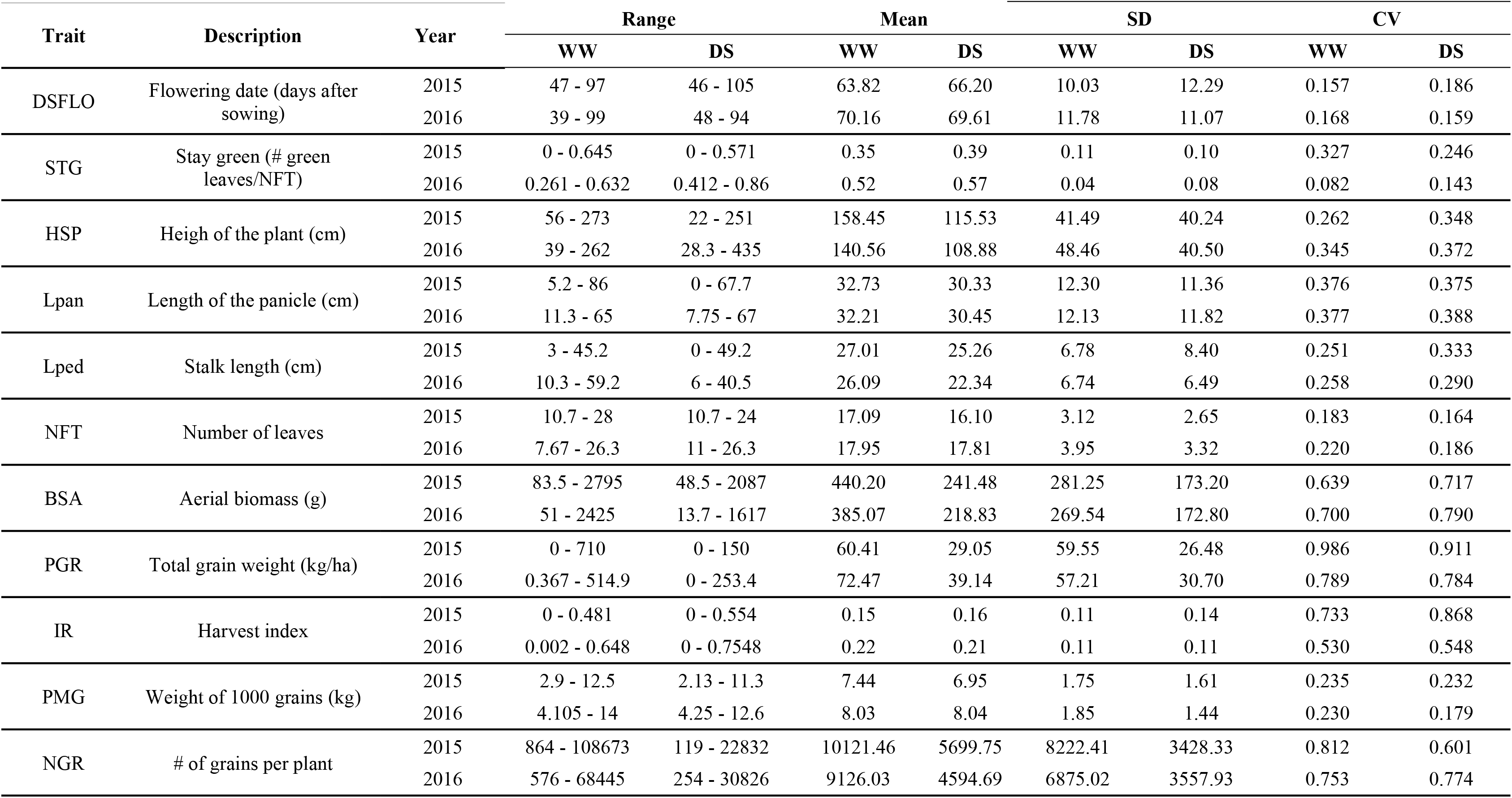
Variations of agro-morphological traits in the pearl millet inbred panel used in this study.

**Table 2.** Analysis of variance and genetic effect on phenotypic variation for STG WW in 2015 and 2016.

|  |  | Df | Sum Sq | Mean Sq | F value | Pr(>F) |
| --- | --- | --- | --- | --- | --- | --- |
| STG WW 2015 | block.unadj | 69 | 1.7305 | 0.0250795 |  |  |
|  | Line | 299 | 4.3200 | 0.0144482 | 11.0845 | <2e-16 *** |
|  | Control | 1 | 0.0007 | 0.0007216 | 0.5536 | 0.4588 |
|  | Control + control.vs.aug. | 298 | 4.3193 | 0.0144943 | 11.1198 | <2e-16 *** |
|  | Residuals | 90 | 0.1173 | 0.0013035 |  |  |
| STG WW 2016 | block.unadj | 69 | 0.24090 | 0.0034913 |  |  |
|  | Line | 288 | 0.56838 | 0.0019735 | 16.493 | <2e-16 *** |
|  | Control | 1 | 0.00000 | 0.0000000 | 0.000 | 1 |
|  | Control + control.vs.aug. | 287 | 0.56838 | 0.0019804 | 16.550 | <2e-16 *** |
|  | Residuals | 73 | 0.00874 | 0.0001197 |  |  |

Significance codes: \*\*\* for 0; \*\* for 0.001; \* for 0.01

**Figure 1.**
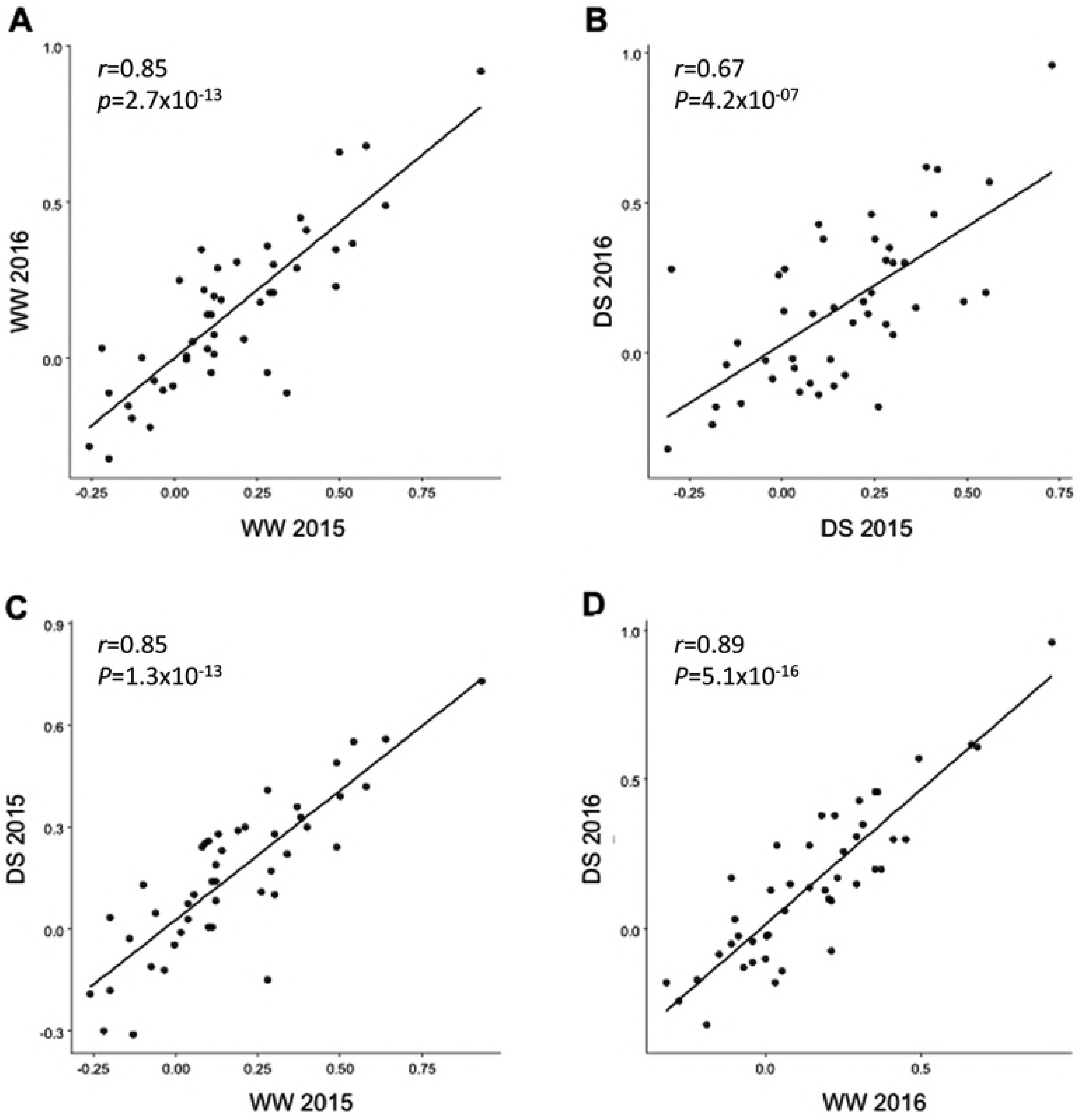
Correlation between agromorphological traits. Plots of the coefficients of correlation calculated between agromorphological traits per pairs. For each graph 45 correlation coefficients associated to 45 pairs of agromorphological traits obtained in different conditions or at different dates were compared among 188 pearl millet lines. The traits were measured in the field in Bambey in 2015 and 2016 with and without hydric stress applied during 30 days at 21 days after sowing A. Plot comparing the coefficients without stress in 2015 and 2016 B. Plot comparing the coefficients with hydric stress in 2015 and 2016. C. Plot comparing the coefficients with and without hydric stress in 2015 D. Plot comparing the coefficients with and without hydric stress in 2016.

**Figure 2.**
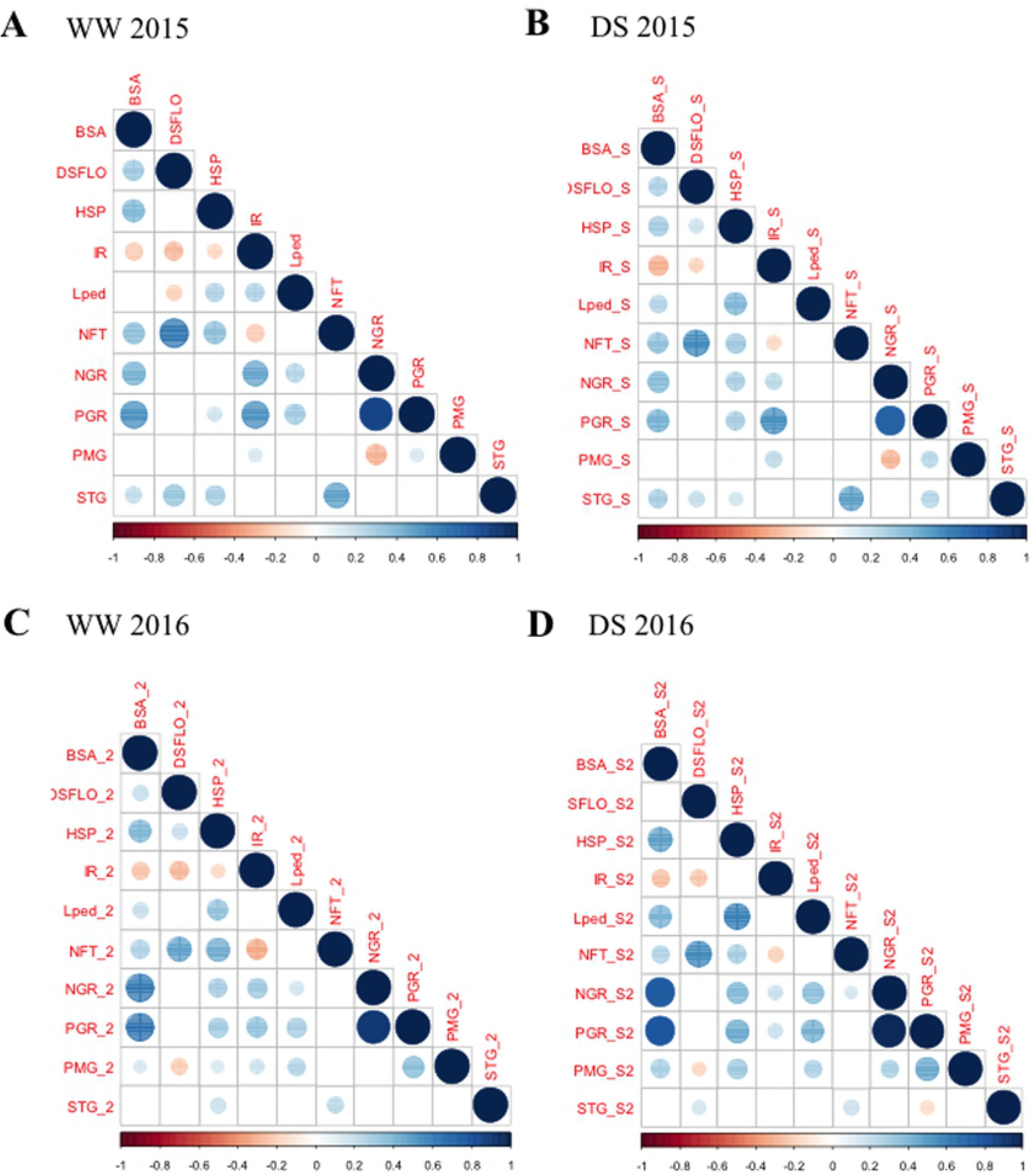
**Correlations between agromorphological traits per pair** measured in the field in Bambey in 2015 and 2016 with and without hydric stress A. No stress in 2015 B. With hydric stress for 30 days at 21 days after sowing in 2015 C. No stress in 2016 D. With hydric stress in 2016. The correlation coefficients are represented by a circle, its size and the strength of its color are proportional to the strength of the correlation. Red and blue indicate negative and positive correlations respectively.

In order to evaluate drought effects on agromorphological traits, values measured under drought stress were divided by values measured under well watered conditions for each trait in both years (Table 3). Drought stress led to a strong and very significant reduction in plant height, above ground biomass, total grain number and weight on both years (Table 3). Panicle length, stalk length and number of leaves were moderately reduced under drought stress. On the contrary, stay green increased slightly but significantly under drought stress on both years. However, the very low harvest index and the failure of some lines to set seeds suggest that the sink demand from grain was low and therefore the observed effect might not be related to a functional stay-green character, i.e. transition from carbon capture to nitrogen remobilization [18]. Flowering date and weight of 1000 grains were unevenly affected on the two years, showing either small or no variation under early drought stress. Harvest index (HI) was not significantly affected by drought on both years. Hence, early drought stress led to a reduction both in biomass and number of grains leading to reduced yield (but not in HI and grain weight). No change in individual grain weight was observed.

**Table 3.** Impact of drought stress on agromorphological traits in pearl millet.

| Trait name | Trait definition | Year | Mean |  | Ratio<br>DS/WW | p-value |
| --- | --- | --- | --- | --- | --- | --- |
|  |  |  | WW | DS |  |  |
| DSFLO | Flowering date (days after sowing) | 2015 | 63.82 | 66.20 | 1.037 | **0.00389** |
|  |  | 2016 | 70.16 | 69.61 | 0.992 | 0.52690 |
| STG | Staygreen (nb green leaves/total number leaves) | 2015 | 0.35 | 0.39 | 1.115 | **1.969E-07** |
|  |  | 2016 | 0.52 | 0.57 | 1.087 | **1.192E-18** |
| HSP | Heigh of the plant | 2015 | 158.45 | 115.53 | 0.729 | **1.699E-41** |
|  |  | 2016 | 140.56 | 108.88 | 0.775 | **5.909E-19** |
| Lpan | Length of the panicle | 2015 | 32.73 | 30.33 | 0.927 | **0.00701** |
|  |  | 2016 | 32.21 | 30.45 | 0.945 | 0.04974 |
| Lped | Stalk length | 2015 | 27.01 | 25.26 | 0.935 | **0.00255** |
|  |  | 2016 | 26.09 | 22.34 | 0.856 | **1.344E-13** |
| NFT | Number of leaves | 2015 | 17.09 | 16.10 | 0.942 | **2.845E-06** |
|  |  | 2016 | 17.95 | 17.81 | 0.992 | **6.154E-01** |
| BSA | Aerial biomass | 2015 | 440.20 | 241.48 | 0.549 | **1.296E-25** |
|  |  | 2016 | 385.07 | 218.83 | 0.568 | **9.547E-21** |
| PGR | Total kernel weight | 2015 | 60.41 | 29.05 | 0.481 | **1.408E-17** |
|  |  | 2016 | 72.47 | 39.14 | 0.540 | **4.877E-20** |
| IR | Harvest index | 2015 | 0.15 | 0.16 | 1.040 | 0.53120 |
|  |  | 2016 | 0.22 | 0.21 | 0.959 | 0.31683 |
| PMG | Weight of 1000 kernels | 2015 | 7.44 | 6.95 | 0.934 | **0.00143** |
|  |  | 2016 | 8.03 | 8.04 | 1.001 | 0.93140 |
| NGR | Number of kernels per plant | 2015 | 10121.46 | 5699.75 | 0.563 | **1.436E-14** |
|  |  | 2016 | 9126.03 | 4594.69 | 0.503 | **3.463E-22** |

Bold numbers correspond to significant differences ( $p < 0.01$ )

The loss in grain production under drought stress conditions was very variable among lines and years. Some lines did not produce grain under drought stress even if their panicle length and aboveground biomass were within the same range as yielding plants. This suggests that drought stress applied at the juvenile phase impacts seed production at later stages. On the contrary some genotypes performed well under drought. Some lines showed consistent behavior across the two years, such as ICML-IS 11194 or ICL-IS 11024 that had a high DS/WW ratio for BSA and PGR in both years. On the other hand, ICML-IS 11066 had low PGR ratios in both years.

Altogether, our data indicate that response to early drought stress was variable in our panel but conserved across lines between the two experiments. This suggests that this population could be used to identify QTLs/genes involved in early drought tolerance through phenotype/genotype association analyses and could a be a source of potential donor lines for drought tolerance.

### Genotypes and population structure

In order to perform genetic association, we generated GBS data on the 188 lines and obtained a total of 3,168,971 unfiltered single nucleotide polymorphisms (SNPs). Filtering on quality led to 392,493 SNPs. The average SNP density was 2.5 per 10 kb.

SNPs were first used to assess the population structure in the panel. The cross-validation error reached a minimum for *K* = 1 suggesting only a single genetic group (S7 Fig). The unweighted neighbor-joining (NJ) tree constructed to illustrate the phylogenetic structure of the panel (Fig 3) displayed also a weak structuration signal. Altogether, this indicates that our panel has a negligible genetic structure.

**Figure 3.**
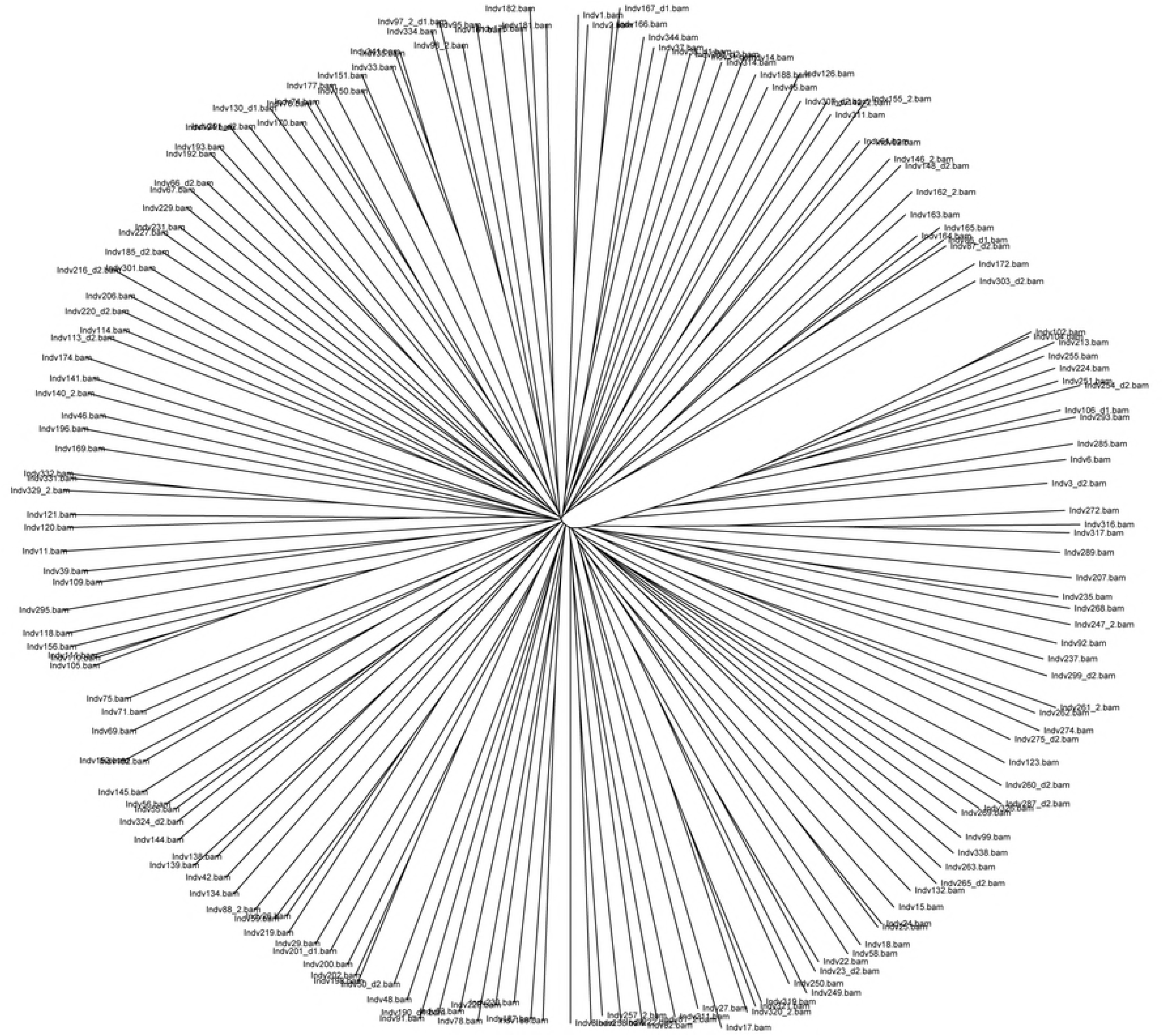
Genetic structure of the inbred panel. NJ tree based on 392 493 SNPs genotyped for 175 pearl millet lines used for GWAS

### Genome wide association study (GWAS)

Genetic association (GWAS) was conducted on 175 inbreds presenting both genetic and phenotypic data. A total of thirteen plants were removed because their genotypic data did not pass filter quality tests. GWAS was performed on all phenotypic traits measured in two years (2015 and 2016) in WW and DS conditions. The most significant associations (*p*−*value* < 10^−5^) detected in the GWAS study are listed in Table 4. The *FDR* for the associations ranged from 0.0001 to 1, with 10 associations having a *FDR* < 0.1.

**Table 4.** GWAS results for the STG and BSA traits.

| Trait | SNP | *p*-value | *FDR* |
| --- | --- | --- | --- |
| STG WW | S6_111262313 | 1.21603721219283E-08 | **0.0011794223317337** |
|  | S6_111262313 | 1.21603721219283E-08 | **0.0011794223317337** |
|  | S6_111262313 | 1.21603721219283E-08 | **0.0011794223317337** |
|  | S6_111262313 | 1.21603721219283E-08 | **0.0011794223317337** |
|  | S2_39226211 | 3.87647802274037E-06 | 0.300780581558053 |
| BSA WW | S4_7055089 | 8.28674626745573E-07 | 0.243010166841778 |
|  | S3_296405038 | 0.0000031432340876036 | 0.243010166841778 |
|  | S3_198858285 | 3.19038839876643E-06 | 0.243010166841778 |
|  | S2_86538637 | 3.91673493887862E-06 | 0.243010166841778 |
|  | S5_145140900 | 4.87502023610317E-06 | 0.243010166841778 |
|  | S5_145140930 | 4.87502023610437E-06 | 0.243010166841778 |
|  | S3_76894947 | 5.46038729608676E-06 | 0.243010166841778 |
|  | S1_49962404 | 0.0000061327667976599 | 0.243010166841778 |
|  | S1_49962418 | 0.0000061327667976599 | 0.243010166841778 |
|  | S3_291115205 | 6.26385896446448E-06 | 0.243010166841778 |
|  | S3_52232262 | 6.90137412656471E-06 | 0.243402681876867 |
| BSA DS | S3_216179591 | 1.39620607372396E-07 | **0.0180478716512176** |
|  | S3_216179589 | 1.39620607372664E-07 | **0.0180478716512176** |
|  | S3_216179592 | 1.39620607372664E-07 | **0.0180478716512176** |
|  | S2_17262004 | 8.76503964281349E-07 | **0.0849750872031571** |
|  | S6_64437341 | 1.15391405443036E-06 | **0.0894954970163208** |
|  | S6_157567763 | 1.39315363532677E-06 | **0.0900420735661673** |
|  | S3_9915484 | 6.92725725514157E-06 | 0.255010678611073 |
|  | S3_13047576 | 7.37303256795503E-06 | 0.255010678611073 |
|  | S3_13047576 | 7.37303256795503E-06 | 0.255010678611073 |
|  | S3_13047576 | 7.37303256795503E-06 | 0.255010678611073 |
|  | S3_13047576 | 7.37303256795503E-06 | 0.255010678611073 |
|  | S5_128247865 | 8.64005390799752E-06 | 0.255010678611073 |
|  | S5_89968400 | 9.51014659847111E-06 | 0.255010678611073 |

Markers with $p < 10^{-5}$ are shown and those with $FDR < 0.1$ appear in bold.

In WW conditions, we identified 4 marker-trait associations (MTAs) for the stay-green character on chromosome 6 (Fig 4, Table 4). These 4 MTAs corresponded to 2 different polymorphisms located at the same position. The stay-green MTA explained 12% of phenotypic variation in the panel in 2015 and a difference of 25% in phenotype was observed between the two alleles (Fig 5). The corresponding polymorphisms were mapped at position 111262313 on chromosome 6 of the pearl millet reference genome [2]. It corresponded to a peak of SNPs/stay green association (Fig 5). Interestingly, this falls within a predicted gene (*Pgl_GLEAN_10013220*) encoding a potential uroporphyrin-III C-methyltransferase (UPM), an enzyme involved in sirohaem and cobalamin biosynthetic pathway. Sirohaem is an essential cofactor involved in inorganic S and N assimilation in plants [19]. No significant association was found in DS conditions but as reported above it might be due to the failure of some lines to set seeds leading to reduced sink demand from grain preventing leaf senescence.

**Figure 4.**
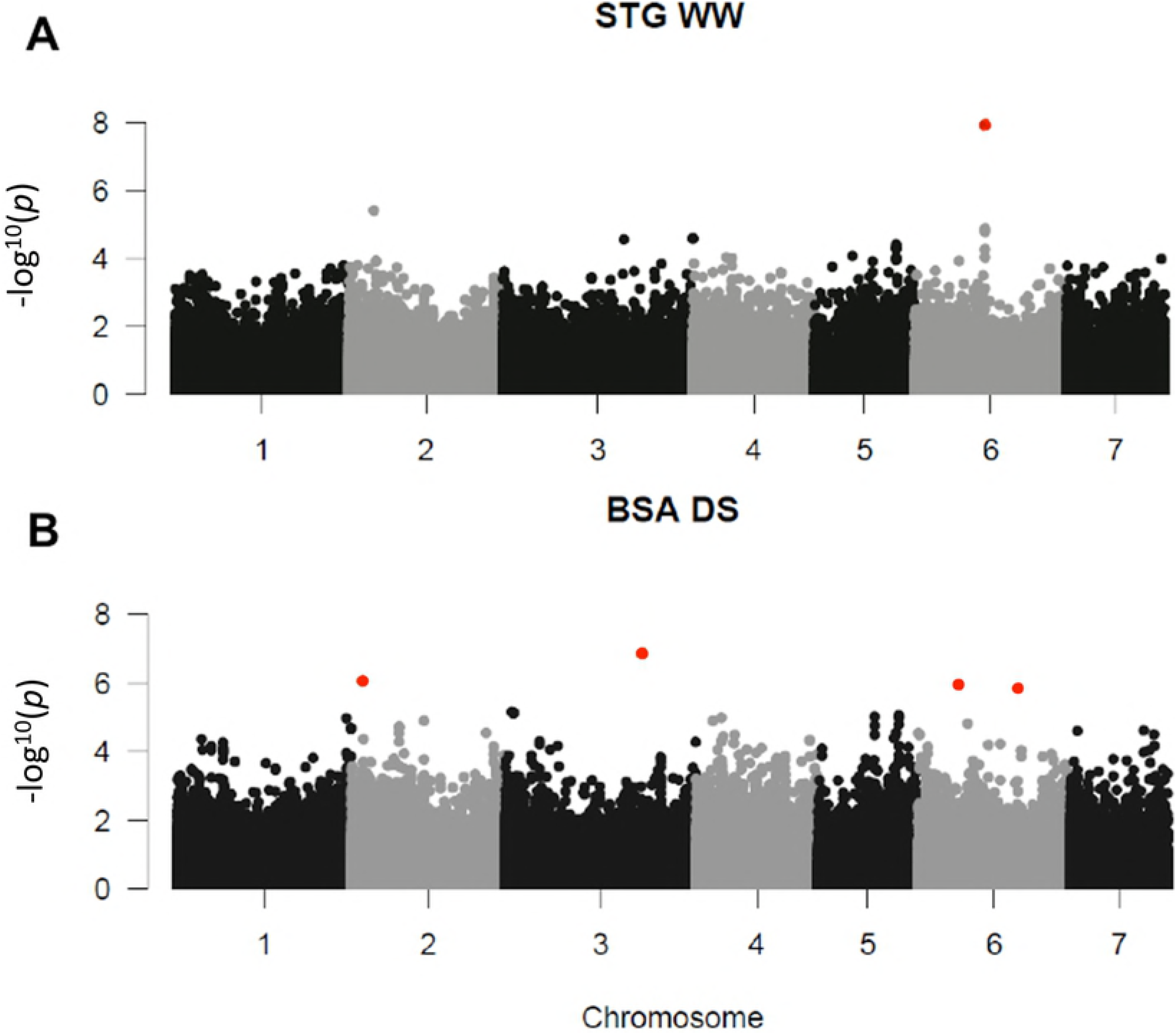
Significant associations for STG and BSA. The whole-genome was scanned using 392 493 SNPs for association with *MAF*>0.05. *p*-values of both years (2015 and 2016) for each trait and each condition were combined using fisher combining probability test. Red dots indicates *FDR* adjusted *p*-value <0.1. A: Manhattan plot for STG in condition WW. B: Manhattan plot for BSA in condition DS.

**Figure 5.**
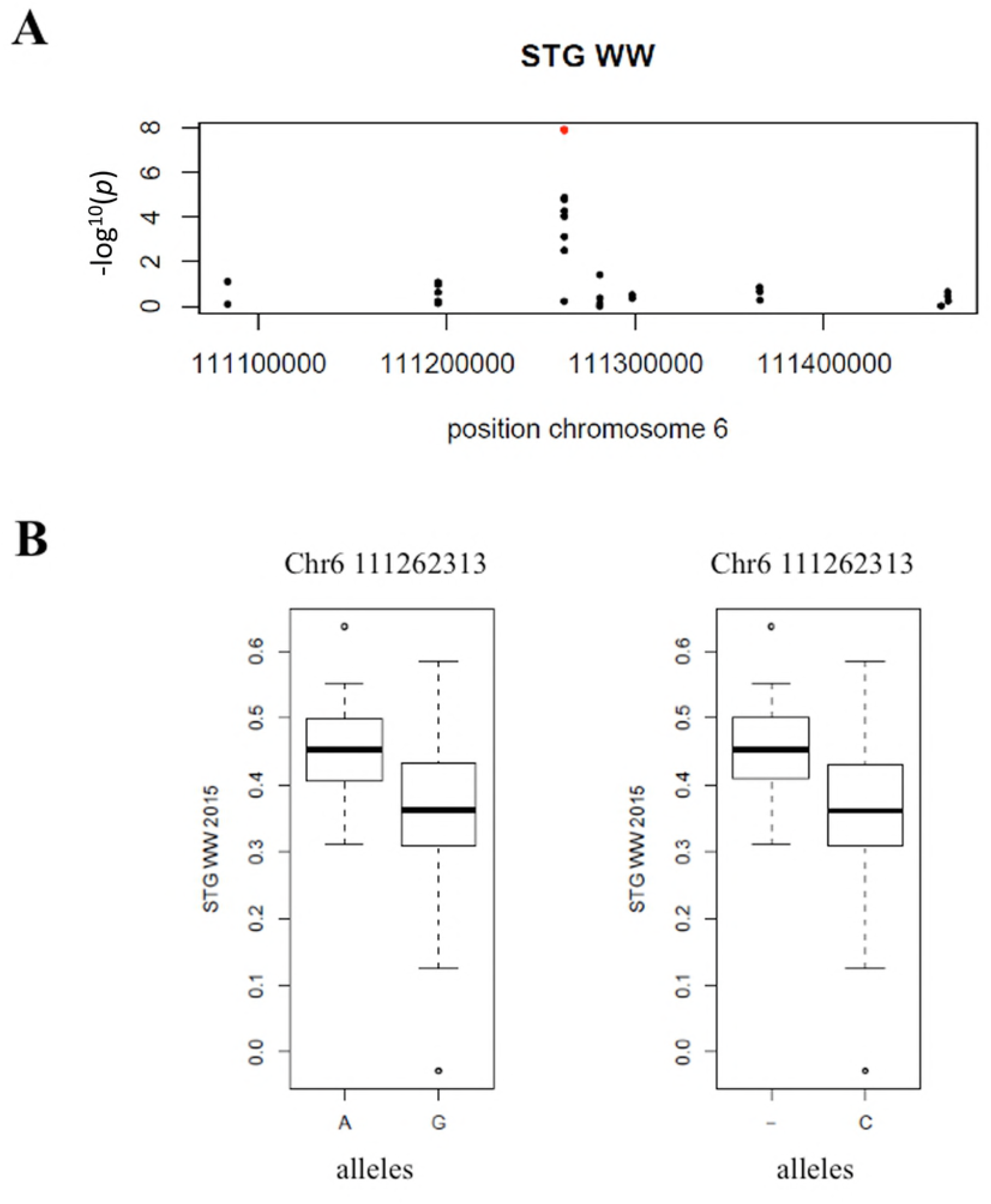
Significant association for STG. A: Zoom in on chromosome 6 position 111,262,313 of the STG WW Manhattan plot. B: Box plots showing phenotypic difference for STG between the two alleles A/G and deletion/C considering one dominant allele respectively “A” and “-“ (deletion) at the significant marker on chromosome 6 position 111,262,313.

Six significant associations were detected for biomass on chromosomes 2, 3 and 6 in DS conditions (Fig 4, Table 4). However, one inbred line had high BSA under DS (Fig 6) and the associations were lost when this line was removed from the analysis. Three of these associated SNPs were very closely located at position 216179589, 216179591 & 216179592 on chromosome 3. These markers explained 19% of the phenotypic variation in the panel. The 3 SNPs were located between two predicted genes *Pgl_GLEAN_10034145* and *Pgl_GLEAN_10034146* encoding a putative 3-ketoacyl-CoA synthase and an unknown protein respectively. 3-ketoacyl-CoA synthases are involved in elongation of C24 fatty acids, an essential condensing step during wax and suberin biosynthesis [20].Comparison of the genome sequence of pearl millet and other cereals revealed an expansion of gene families involved in wax and suberin biosynthesis that was proposed to have contributed to pearl millet adaptation to drought and heat stress [2]. Accordingly, changes in wax and/or suberin biosynthesis linked to a 3-ketoacyl-CoA synthase encoding gene might be responsible for quantitative changes in biomass under drought stress conditions as observed in our field trials.

**Figure 6.**
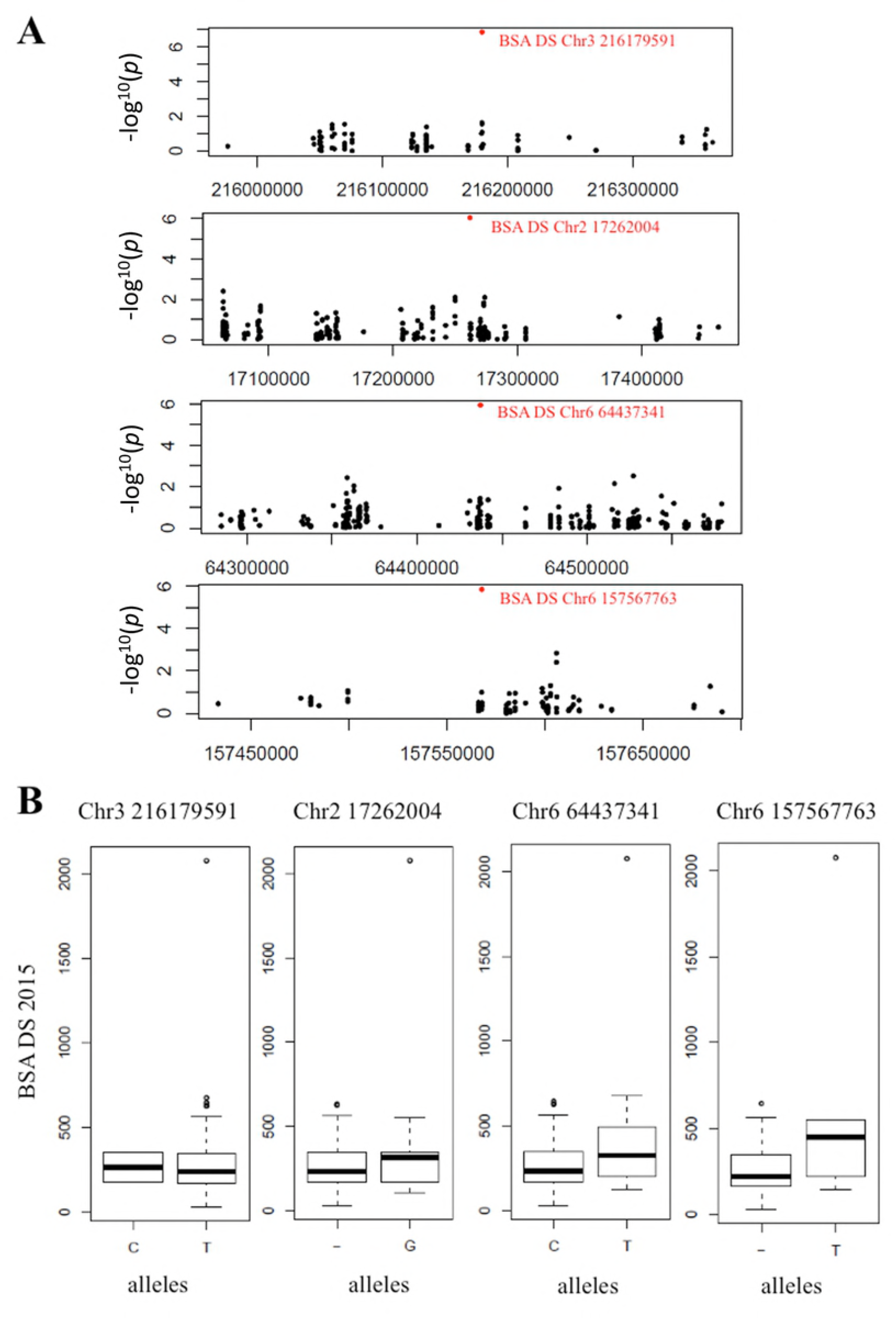
Significant associations for BSA under drought stress. A: Manhattan plots, zoom in on chromosome 2, 3 and 6 for each significant association of BSA DS. B: Box plots showing phenotypic difference for BSA between the two alleles at these positions: 216179591 of chromosome 3, 17262004 of chromosome 2, 64437341 and 157567763 of chromosome 6, considering the allele at lowest frequency as dominant.

The association on chromosome 2 was mapped at position 17262004. It explained 14% of the phenotypic variation in the panel (Figure 6). The SNP is located in the predicted gene *Pgl_GLEAN_10005405* that encodes a putative chloroplastic threonine dehydratase. This enzyme catalyzes the formation of alpha-ketobutyrate from threonine. This catabolic reaction might change the amino acid metabolism in response to stress and facilitate photorespiration. Photorespiration is known to sustain CO_2_ fixation capacity in conditions of water stress and to improve drought tolerance in cereals [21].

Two SNPs on chromosome 6 were significantly associated with biomass production in early drought stress conditions. The first one was mapped at position 64437341, between two predicted genes *Pgl_GLEAN_10037360* and *Pgl_GLEAN_10037359* encoding an unknown protein and a SPA1-related 3 protein homolog. The SPA proteins contain serine/threonine kinase-like and WD40 protein domains and are involved in signal transduction during photomorphogenesis [22]. It has been shown that SPA1 modulates MYC2-mediated ABA and JA responses [23] that are known regulators of plant response to stresses including drought. This association explained 14% of the phenotypic variation in the panel (Fig 6). The second SNP associated with biomass production under drought was located at position 157567763 between two predicted genes *Pgl_GLEAN_10036945* and *Pgl_GLEAN_10036946* encoding an unknown protein and a putative E3 SUMO-protein ligase respectively. E3 SUMO-protein ligases regulate protein sumoylation and have been involved in tolerance to a number of abiotic stresses including drought [24]. In Arabidopsis, E3 SUMO-protein ligase SIZ1 contributes to the accumulation of SUMO-protein conjugates induced by drought stress [24]. Moreover, SIZ1 was demonstrated to regulate growth and drought stress response [24] as well as abscisic acid signaling [25].

## Discussion

As little is known about the impact of early drought episodes in pearl millet, we analyzed the response of a panel of 188 pearl millet inbred lines to drought stress during the juvenile phase for 2 years in field conditions in Senegal. Drought stress was applied from 21 days after germination (DAG) to 49 DAG. On drawback of such approach is that while the stress was applied at the same fixed time for all lines, we have a large diversity of flowering time in our panel and all plants did not therefore face stress at the exact same developmental stage. However, the earliest flowering genotypes in our panel flowered at 47 or 39 DAG in well watered environment and at 46 or 48 DAG in drought environment depending on the year. Beginning of flowering therefore corresponded almost to the end of drought stress for the early flowering lines. The mean flowering time in our panel was 66 and 69 for the two years respectively for drought stress treatment, so between 17 to 20 days after the end of the stress.

Consequently, the majority of the lines flowered after drought stress. Accordingly, we did not find any significant relation between flowering time and PGR (grain yield) or panicle length under stress in both years. One strategy to avoid such issue would have been to work with groups of lines having similar flowering time but this would have limited the number of lines available for analysis and therefore the potential genetic diversity usable for gene discovery.

We found that early drought stress led to a reduction in both grain and biomass production. Limited change in grain weight (as would be expected for a terminal drought stress) was observed only one year out of 2 suggesting that most pearl millet lines were able to adapt their biomass to water availability in order to produce a limited number of viable seeds. Yield loss in early drought stress conditions was very variable among lines. Some lines did not produce grain under drought stress even if their panicle length and aboveground biomass were within the same range as yielding plants. On the other hand, some lines did perform better in drought stress conditions. Nevertheless, our results clearly show that drought episodes during the vegetative phase of the plants can have a dramatic impact on yield even if water is available during the end of the cycle. By reducing water availability at this early stage, plants were facing drought stress during the juvenile phase, when the young panicle develops, deep inside the meristem. How this impacts seed formation later in the cycle when water supply remains to be deciphered.

We exploited the phenotypic diversity observed in our panel to perform GWAS to uncover the genetic bases of tolerance to early drought stress in pearl millet. We identified 10 SNPs associated with stay-green and biomass phenotypes corresponding to 5 potential QTLs. As pearl millet biomass is increasingly used as fodder, biomass quantity and quality (of which stay-green is an important character) are becoming important targets for breeding. Moreover, changing temperature and photoperiod associated to climate change are known to impact senescence [26]. Regulation of stay-green is therefore important to design varieties better adapted to future climate in West Africa. Stay-green is linked to the remobilization of N from leaves for grain filling leading to leaf senescence [18]. Slower or inhibited remobilization leads to stay-green and while it is linked with drought stress tolerance in some crops (sorghum for example) this is not always the case [18]. Interestingly, the SNPs associated to stay-green are located in a gene encoding a potential uroporphyrin-III C-methyltransferase, an enzyme of the sirohaem and cobalamin biosynthetic pathway [27]. Sirohaem is an essential component of the plant sulphite and nitrite reductase enzymes [27], [28]. It has been proposed that enhanced N uptake might contribute to stay-green and drought tolerance in sorghum [29]. Accordingly, increased nitrite reductase activity in tobacco leads to a stay-green phenotype [30]. We can hypothesize that polymorphisms in the UPM gene might impact sirohaem biosynthesis and as a consequence nitrite reductase activity leading to increased nitrogen assimilation and potentially to increased stay green. Further work will be required to test this hypothesis. No significant association was found for stay green in DS conditions but it might be due to the failure of some lines to set seeds leading to reduced sink demand from grain preventing leaf senescence. Interestingly, the harvest index of the germplasm we tested was also relatively low (around 0.15), suggesting that the sink strength of the panicles was limited in comparison to standard elite genotypes. This suggests that the stay-green we measured is not an indication of a drought stress tolerance but rather an indication of the balance of the remobilization process between the reproductive and vegetative organs.

Pearl millet is a tough crop that can survive and yield in harsh low water and high-temperature conditions [2], [3], [8]. As such it is a great model to identify mechanisms involved in drought and heat stress tolerance in cereals. Here we identified 4 loci associated with increased biomass production under early drought stress. These associations were dependent on the inclusion of one line with a strong BSA phenotype under drought stress so they will need to be confirmed. Genes encoding proteins involved in signal transduction pathways related to drought were found to co-localize with 2 of these loci while a third associated SNP was found in a gene encoding an enzyme involved in threonine catabolism. The last loci associated with increased biomass under drought stress contained a gene encoding an enzyme (3-ketoacyl-CoA synthase or KCS) that catalyzes the elongation of C24 fatty acids during both wax and suberin biosynthesis [20]. Cuticular waxes are the main transpiration barriers in leaves and a correlation between wax content and drought tolerance has been reported in many crops [31]. Suberin is a polymer made of aliphatic and aromatic compounds and confers hydrophobic characteristic to the walls of certain root cells (in particular in the endodermis), providing barrier properties against water diffusion [32]. It has been associated with drought tolerance in some species. For instance, the Arabidopsis *enhanced suberin 1* (*esb1*) mutant, characterized by twofold increased root suberin content, has increased water-use efficiency and drought tolerance [33], [34]. Suberin deposits have been observed in the endodermis and exodermis in pearl millet roots [35]. Interestingly, the gene families related to wax, cutin and suberin biosynthesis and transport are expanded in pearl millet compared to other cereal crops [2]. It has been proposed that this might have contributed to heat and drought tolerance in pearl millet [2], [3]. The KCS gene in the locus associated to biomass production under early drought stress might be related to this. The material we identified in this study will be instrumental to test this hypothesis and it will be interesting to analyze in future studies the relation between drought tolerance and/or suberin composition and content in lines contrasted for drought tolerance.

In conclusion, our results reveal new potential mechanisms regulating response to drought stress and the stay green trait in pearl millet and open the way to use genomic selection to breed pearl millet lines with improved yield and drought stress tolerance.

## Acknowledgments

This work was supported by the IRD, the French Ministry for Research and Higher Education (PhD grant to SP) and the NewPearl grant in the frame of the CERES initiative by the Agropolis Fondation (N° AF 1301-015 to LL, as part of the “Investissement d’avenir” ANR-l0-LABX-0001-0l) under the frame of I-SITE MUSE (ANR-16-IDEX-0006) and by the Fondazione Cariplo (N° FC 2013-0891).

## Supporting information

**S1 Fig. Field experimental set up (2015).** Plants were organized in 70 blocks with 20 plants per block along 18 rows and 24 columns. A control inbred line (yellow) was installed regularly to correct for soil heterogenity in the field.

**S2 Fig. Field experimental set up (2016).** Plants were organized in 70 blocks with 20 plants per block along 18 rows and 24 columns. A control inbred line (yellow) was installed regularly to correct for soil heterogenity in the field.

**S3 Fig. Exemple of spatial correction of Lped for the DS trial of 2016 with SpATS.** A: Original data (stalk length in cm) plotted against row and column positions in the field. Plots corresponding to control line appear in diagonal, according to field experimental setup (SupFig1 & 2). B: Value estimated by the model, taking into account spatial and genetic effect. C: Residuals of the model, spatially independent. D: Fitted spatial trend. E: Genotypic BLUEs (best linear unbiased estimates) F: Histogram of BLUEs

**S4 Fig. Fraction of transpirable soil water (FTSW) in the well-watered (WW) and drought stress (DS) treatments in 2015 and 2016.** Volumetric soil moisture (VSM) was monitored along the soil profile (0-60 cm and 60-120 cm) at different locations inside the field by frequency domain reflectrometry through PVC pipes (Diviner 2000, Sentek Sensor Technologies, SA, Australia) and converted as percent of FTSW as follow: (VSMact – VSMmin) / (VSMmax – VSMmin) where VSMact is the actual averaged VSM values measured every 10 cm between 0-60 and 60-120 cm, VSMmax is the maximum VSM value observed in the WW treatment and VSMmin is the minimum VSM value observed in the DS treatment. The dashed red line represents the FTSW of 40% below which the water is considered as limiting for pearl millet growth. Points represent mean (n=4-10) ± se.

**S5 Fig. Changes in canopy temperature between well-watered (WW) and drought stress (DS) treatments in 2015 and 2016.** Canopy temperature (CT) was measured on a sunny day (at 51 DAS in 2015 and 2016) typically between 11AM and 12PM on two leaves per plot using an infrared sensor (Company…). Averaged CT values for all genotypes within one treatment of one experiment were used to prepare boxplot. ***: p < 0.001 using ANOVA statistical test.

**S6 Fig. Distribution of agromorphological characters for 2015 WW (A), 2015 DS (B), 2016 WW (C) and 2016 DS (D)**

**S7 Fig. Graph of Cross-Entropy calculated for population structure from one to 20 groups.**

**S1 Data. Corrected field trials data.**

